# Host cell remodeling via cyclin dependent kinases drives Ebola virus replication and transcription

**DOI:** 10.64898/2026.03.25.714206

**Authors:** Tatiana M. Shamorkina, Danni Snikkers, Albert J. R. Heck, Joost Snijder

## Abstract

Ebola virus (EBOV) causes re-emerging outbreaks of viral hemorrhagic fever with high case-fatality rates. As a negative-sense single stranded RNA virus, EBOV depends on its RNA-dependent RNA polymerase (L protein) to transcribe and replicate the viral genome. This takes place in cytoplasmic inclusion bodies that form following host cell remodeling involving yet unknown signaling pathways. Here, using mass spectrometry-based (phospho-)proteomics, we profiled global protein abundance and site-specific phosphorylation in HEK293T cells expressing an EBOV minigenome system. Our data reveal EBOV-induced rewiring of the host proteome and phosphorylation signaling landscape, including perturbations in cell cycle control, cytoskeletal organization, innate immune regulation, and DNA damage response. Kinase network analysis revealed that Hippo pathway kinases and especially cyclin dependent kinases like CDK2 are central drivers of EBOV replication and transcription. The functional necessity of these signaling pathways is demonstrated via inhibition of CDK family kinases with small molecule inhibitors, which halted EBOV minigenome replication and transcription when administered in the low micromolar range, demonstrating that these pathways represent putative antiviral targets.

## Introduction

Since the initial discovery of Ebola virus (EBOV) in 1976, several members of its genus *Orthoebolavirus* have caused recurring outbreaks of viral hemorrhagic fever in Central and West Africa (1,2). Most recently, the Public Health Ministry of the Democratic Republic of the Congo declared an outbreak of Ebola Virus Disease (EVD) in September 2025 (3). This outbreak involved 64 confirmed and probable cases, including 45 deaths (4). This represented the 35^th^ outbreak on record, including also outbreaks of the related Sudan virus, Bundibugyo virus, and Tai Forest virus (2,5). Outbreaks of EVD are characterized by their high case fatality rates of up to 90% and typically range between 10-1000 diagnosed cases (1,6–8). Starting in December of 2013, the historically largest outbreak of EVD occurred in West Africa, causing an estimated 28,610 cases, resulting into 11,308 deaths (9). The outbreak was concentrated in Sierra Leone, Guinea and Liberia, but spread as far as Mali, Senegal and Nigeria, as well as the United States of America, the United Kingdom, Spain, and Italy (primarily by infected travelers and active health care workers returning from duty) (9). While an effective EBOV vaccine was first tested during the West African epidemic and later also employed in following outbreaks (5,10), no proven vaccines against other members of *Orthoebolavirus* are available, and EVD treatments have shown only limited success with no effective cure available (5).

*Orthoebolavirus* is a genus within the family *Filoviridae* (11). *Orthoebolaviruses* are negative sense single stranded RNA viruses with a genome of ca. 18 kb encoding seven genes that are responsible for viral replication, transcription, assembly and budding (6). The longest of its genes, L, encodes the viral RNA-dependent RNA polymerase (RdRp), responsible for genome replication, transcription, and mRNA capping (1,6). Together with transcription regulator VP30, co-factor VP35, and the nucleoprotein NP, this minimal set of genes is sufficient for active viral genome replication and transcription in human cells (1,6,7). These proteins associate with viral genomic or messenger RNA to form ribonucleoprotein complexes (RNPs). The remaining three genes of the ebolavirus genome encode VP24, which complements NP for assembly of the viral genome into tightly wound helical filaments; the matrix protein VP40 which sits on the luminal side of the viral envelope and recruits assembled nucleocapsids to budding sites at the plasma membrane; and the viral glycoprotein GP, which sits in the viral envelope to mediate cell attachment, host receptor NPC1 binding, and entry through its fusion activity of the viral envelope with the host cell membrane (1,6,7).

Replication and transcription of EBOV occur in cytoplasmic inclusion bodies (12). Owing to its negative-sense genome, the viral RNA cannot serve directly as a template for protein synthesis and instead requires the RNA-dependent RNA polymerase L to generate positive-sense mRNAs that initiate translation by host ribosomes (12). Consequently, the polymerase complex must be co-packaged into budding virions to enable transcription upon entry into newly infected cells (6). The mechanisms by which EBOV proteins subsequently rewire the host cell to promote formation of the cytoplasmic inclusion bodies, recently also referred to as viral factories, remain unclear. Intrinsic biophysical properties of the EBOV genome and proteins likely contribute to the formation of these biomolecular condensates in the cytoplasm (6,12). However, previous studies have revealed an extensive interaction network with host proteins, along with clear evidence that host-mediated phosphorylation and other post-translational modifications promote immune evasion and remodel the cellular environment to facilitate EBOV replication and gene expression (13,14).

As a class 4 biological agent, research on EBOV relies on appropriate model systems for reasons of biosafety and -containment. Because L, VP35, VP30 and NP are together sufficient for active genome replication and transcription, their co-expression with viral genomic RNA in human cells induces the formation of RNPs in cytoplasmic inclusions, resembling those observed during authentic infection (12,15,16). This led to the development of minigenome systems, which have been extensively used for the past three decades as models to study filovirus replication, transcription, and the associated host cell remodeling and suppression of innate immune signaling (15,17). In the most common and minimal minigenome implementation, separate plasmids encoding L, VP35, VP30, and NP are co-transfected with a minigenome plasmid, which encodes a reporter gene (commonly GFP or luciferase), flanked by viral UTR’s, transcription start/stop sites, and the genomic leader and trailer, thus producing a functional minigenome that serves as a substrate for L activity and formation of RNPs in cytoplasmic inclusion bodies (15).

Previous studies of proteome-wide ebolavirus-host interactions have focused primarily on protein-protein interactions using affinity purification or proximity labeling of individually expressed viral proteins or the abovementioned minigenome systems (6,13,18–20). Such studies have identified a multitude of associated pathways that are functionally involved in ebolavirus replication and transcription, including cell cycle regulation, cytoskeletal remodeling, RNA metabolism, protein folding, and translation. The host factors involved include the proteins RBBP6, hnRNP L, hnRNPUL1, and PEG10, which interact with VP30 through a PPxPxY motif, mimicking the VP30-NP interaction, to modulate transcription activity of L or conversely act as restriction factors that limit viral RNA synthesis (14,20). Host proteins GSTP1 and UPF1, involved in nonsense-mediated mRNA decay, have similarly been identified to interact with L, with their interaction promoting viral RNA synthesis. Similarly, VP35 interacts with the mRNA decapping complex through the scaffold subunit EDC4 (19), and dynein light chain LC8 is known to interact with the VP35 N-terminal oligomerization domain (21), both interactions promoting viral RNA synthesis. Additionally, host protein STAU1, RUVBL1, SMYD3, and CAD have been identified to interact with ebolavirus proteins to promote RNP formation, increase RNA metabolism and thereby support viral RNA synthesis (6,22–24).

Host cell remodeling by ebolavirus also occurs through post-translational modifications of viral and host proteins, including through ubiquitination, SUMOylation, hypusination, and most notably phosphorylation-based signaling that regulates the replication/transcription activity of L and suppresses innate immune signaling (6). For instance, dynamic (de-)phosphorylation of the VP30 N-terminus at serine 29 by host kinases SRPK1/SRPK2 or protein phosphatases PP2A-B56 and PP1 is known to regulate the switch between replication and transcription activity of L, with phosphorylation at S29 promoting replication (25–27). The same residue can also be modified by LATS1/2 kinases, part of the Hippo signaling pathway, which further modulates ebolavirus egress through VP40 phosphorylation (28). Suppression of interferon type I (IFN-I) signaling is another major part of the host cell remodeling that takes place to facilitate ebolavirus replication and transcription. Most notably, VP35 inhibits IRF-3 by competitively interacting with kinases IKKε and TBK-1, as well as by sequestering STING to the cytoplasmic inclusions to suppress the interferon response (29). Similarly, VP35 inhibits PKR autophosphorylation and activation, to prevent eIF2α phosphorylation and allow for continued translation and viral protein production (30).

While ebolavirus replication and transcription thus require extensive host cell remodeling for RNP formation in viral factories and suppress innate immune signaling (6), no comprehensive proteome-wide investigation into the associated changes in host cell signaling by phosphorylation has yet been reported. Here we describe (phospho-)proteomics experiments to monitor the proteome-wide changes in cells that facilitate ebolavirus replication and transcription, using human HEK293T as host cells for an active ebolavirus minigenome system. We report a moderate shift in the proteome, dominated by the expression of the viral proteins of the minigenome system, along with significant changes in about 40 host proteins, including known restriction factors such as TRIM14 and ZDHHC20 (31,32). In parallel, we observed extensive alterations in phosphorylation signaling, with over 300 sites showing significant changes in the minigenome transfected cells vs. negative control. These included increased site-specific phosphorylation of RBBP6, C1QBP and CD300a, known EBOV interactors or inhibitors of innate immune signaling via the interferon response. Furthermore, we identified 23 phosphorylation sites on the viral RNP components NP, VP30 and VP35, including several previously unreported sites. Pathway analysis revealed significant changes in cell cycle regulation, RNA metabolism, innate immune signaling, cytoskeletal remodeling and DNA-damage response (DDR) signaling. Motif analysis highlighted CDK2, ATM/ATR (DDR), and PRKDC2 as central kinases regulating the response. Disruption of these phosphorylation signaling pathways with small molecule inhibitors (at micromolar concentrations) of especially the cyclin dependent kinases shut down EBOV replication and transcription in the minigenome systems, confirming their functional significance. These observations provide a systems level overview of phosphorylation-based signaling involved in host cell remodeling upon ebolavirus replication and transcription, highlighting a central role for cell cycle remodeling. These identified host signaling pathways may provide promising targets for the development of broad spectrum *Orthoebolavirus* inhibitors.

## Results

### Host proteome and phosphoproteome remodeling is driven by EBOV replication and transcription

To investigate how active EBOV replication/transcription impacts the host proteome and phosphorylation-based cell signaling, we used a well-established EBOV minigenome assay (15) for global proteomic and phosphoproteomic profiling of transfected human HEK293T cells (Figure 1A). Transfection with EBOV L, VP30, VP35, NP, and minigenome (with eGFP reporter) plasmids led to notable eGFP expression within 24 hours, indicative of active minigenome replication and transcription (Figure S1A). Cells were collected in biological triplicates at 6 and 24 hours post transfection. The 6-hour timepoint was initially selected to capture early phosphorylation events in the transfection system. However, at this time point, only NP, VP30, and VP35 were detected, whereas the EBOV L polymerase was not yet observed (Figure S1B). For the 24-hour timepoint, all viral proteins were detected with NP, VP35 and VP30 being amongst the most abundant proteins in the cell (Figure 1B). Based on these findings all subsequent analyses were focused on the 24-hour post transfection timepoint.

**Figure 1.**
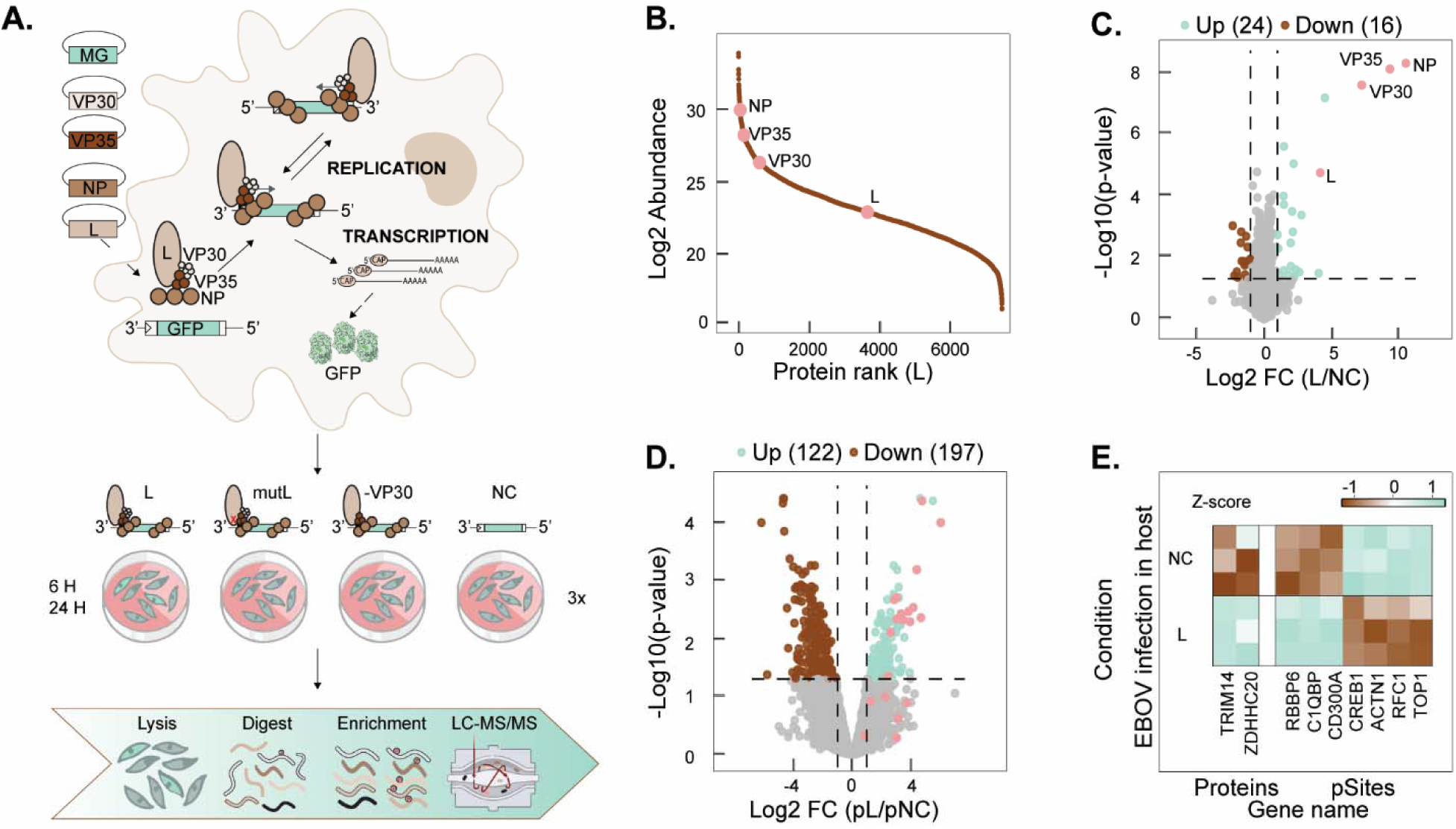
Global proteomics and phosphoproteomics analyses of human cells expressing an Ebola virus (EBOV) minigenome model system of transcription/replication. **(A)** Schematic overview of the minigenome system, tested conditions and workflow. HEK293T cells were transfected with EBOV RNP plasmids encoding L, NP, VP35, VP30 proteins and a eGFP-reporter minigenome (MG). Following transcription by host RNA polymerase II, viral RNA is replicated and transcribed by the RNP complex, with eGFP expression serving as a readout of replication/transcription activity. Four conditions were analyzed: L (functional ribonuclear complex), mutL (inactive polymerase L with N743A mutation in catalytic domain), −VP30, and NC (MG only). Cells were collected at 6 and 24 hrs post-transfection, followed by proteome and phosphoproteome analysis by LC-MS/MS. (B) Protein rank plot of L condition shows the EBOV RNP proteins (pink) within the proteome rank. (C,D) Volcano plots represent Log2-transformed abundance fold change (FC) of L condition versus NC 24 H post-transfection for protein groups (C) and phosphosites (D). Mint green represents the increase in abundance of host proteins in L condition (FC > 1, p-value <0.05). Brown represents the decrease in abundance of host proteins in L condition (FC < -1, p-value <0.05). Pink highlights EBOV proteins or EBOV protein phosphosites. Overall, 7447 protein groups and 10126 phosphosites could be quantified. (E) Regulated proteins and phosphosites that are either previously associated with EBOV infection or key components of enriched WikiPathway “EBOV infection in host” (p-value<0.05). The abundances were standardized using Z-scoring.

We conducted the experiments with a range of transfection controls to delineate the effects of transfection, polymerase activity, and the contributions of replication or transcription (Figure 1A). First, all the minigenome assay plasmids were transfected in HEK293T cells, representing fully active EBOV replication and transcription (condition denoted ‘L’). The ‘L’ condition was repeated using the same plasmids except exchanging to an inactive L protein harboring a point mutation N743A in the catalytic pocket of its polymerase domain, representing inactive but fully assembled EBOV RNP complexes in the cell (denoted ‘mutL’). Furthermore, we included a transfection with all minigenome plasmids except the transcription factor VP30, representing active EBOV replication only, without transcription (denoted ‘-VP30’). Finally, for the negative control condition, the cells were transfected with the equivalent amount of the eGFP reporter minigenome plasmid only (denoted ‘NC’). We used mass spectrometry-based proteomics to quantify system-wide changes in protein abundances and site-specific phosphorylation between the conditions listed above. Principal component analyses showed a separation only of the ‘L’ condition from all others at the proteome level (Figure S1C), whereas phosphorylation patterns were completely separated in PC1 and PC2 directions for all tested conditions, with deficient mutL and -VP30 conditions still in closest proximity to each other (Figure S1D). The quantitation in proteome and phosphoproteome was found to be consistent between biological replicates (Figures S1E-F).

In total 7447 protein groups and 10126 class I (>75% localization probability) phosphorylation sites that belonged to 3177 protein groups were quantified across all conditions (Figure S1G and S1H, Suppl data 1). The number of protein groups showing significant differential abundance relative to NC was comparable across conditions (ca. 40), indicating only a moderate shift at the proteome level (Figure S1I), with the transfected EBOV proteins exhibiting the largest fold changes (Figure 1C; Figure S1K). Given the small number of proteins with significant change in abundance, the gene ontology enrichment analysis lacked sufficient statistical power and therefore did not return any enriched terms. However, amongst the significantly changing proteins TRIM14, ZDHHC20, SOX2, FLCN, and ATXN1L have been previously implicated in the host cell response to EBOV and other viruses (31–35) (Figure 1E, Suppl Data 1). The single biggest change in host protein expression was observed for WTIP, an adaptor protein involved in Hippo signaling (36).

Whereas the changes in protein abundance in response to EBOV minigenome activity were modest, phosphorylation dynamics exhibited a major shift with 319 phosphorylation sites of 265 proteins being differentially regulated in cells with an actively replicating and transcribing minigenome system (Figures 1D, S1J and S1L). None of the host proteins with significant changes in their phosphorylation sites showed significant changes in protein abundance (Figure S1M), underscoring that the observed changes in phosphorylation were not a byproduct of altered protein expression, but rather due to active phosphorylation signaling as a central early host cell response during the EBOV replication cycle.

As for the inactive EBOV RNP complex (mutL) and EBOV replication only (-VP30) systems, much fewer changes in phosphorylation were detected, with most regulated sites located on EBOV proteins (Figure S1J and S1L). The remaining regulated phoshosites (PRPF4 S34, UBXN4 S180, POLR1G S309) were associated with protein metabolism, suggesting aberrant protein production in the mutL condition characterized by transfection of the inactive protein complex. Meanwhile, in the ‘-VP30’ condition, phosphorylation changes (e.g., SFSWAP S834, BCLAF1 S181, LRRFIP1 S471, QSER1 T1341, PARP1 T373, CCAR2 T454, and TAF15 S226) were predominantly associated with transcriptional regulation, RNA processing, and RNA surveillance, suggesting engagement of host gene-expression and RNA metabolism pathways in response active viral replication. Furthermore, the overlap between differentially abundant proteins and phosphosites in the L condition versus mutL and ‘-VP30’ was limited (Figures S1N, S1O, S3A). These findings indicate that host cell remodeling on the proteome and phosphoproteome levels is driven for the most part by the full range of L activity, including both replication and transcription. Therefore, we focused on the changes in ‘L’ vs ‘NC’ in the more detailed analyses described below.

Initial pathway enrichment analysis of the regulated phosphorylation sites in the EBOV replicating and transcribing cells (L) returned the pathway “EBOV infection in host” with a subset of the phosphosites and their corresponding proteins (Figure 1E). One of these proteins, RBBP6, was previously shown to directly interact with EBOV VP30 and negatively regulate EBOV replication (14), but the involvement of phosphorylation signaling in this process has not been previously reported. These observations, together with the upregulation of proteins previously implicated in antiviral responses (TRIM14 and ZDHHC20) (31,37), support the notion that the EBOV minigenome system representa a realistic model of EBOV replication and transcription, indicating that a considerable part of the host cell remodeling during EBOV infection is driven by the EBOV replication and transcription machinery itself.

### Phosphorylation of the EBOV ribonuclear protein complex by host kinases

Amongst the 319 significantly changing phosphorylation sites, 23 mapped to the transfected EBOV proteins NP, VP35 and VP30, but none to L (Figure 2). These include the previously established phosphorylation sites S29/S31 on VP30 (responsible for switching between replication and transcription activities) (38), as well as sites S187 and S317 of VP35 (shown to regulate EBOV replication) (39). Most of the EBOV phosphorylation site sequence motifs didn’t predict any specific kinase responsible for the phosphorylation (Figure 2). However, VP30 S283 as well as NP S581, T631, and S691 were predicted to match ATM and DNAPK kinase recognition motifs (Figure 2, Table S1), which are both involved in the DNA damage response (DDR) pathway. Furthermore, CKII kinase motifs were predicted for NP S587, S644, and S691, along with two singular motifs for PKA (VP35 S205) and PKC (NP S413).

**Figure 2.**
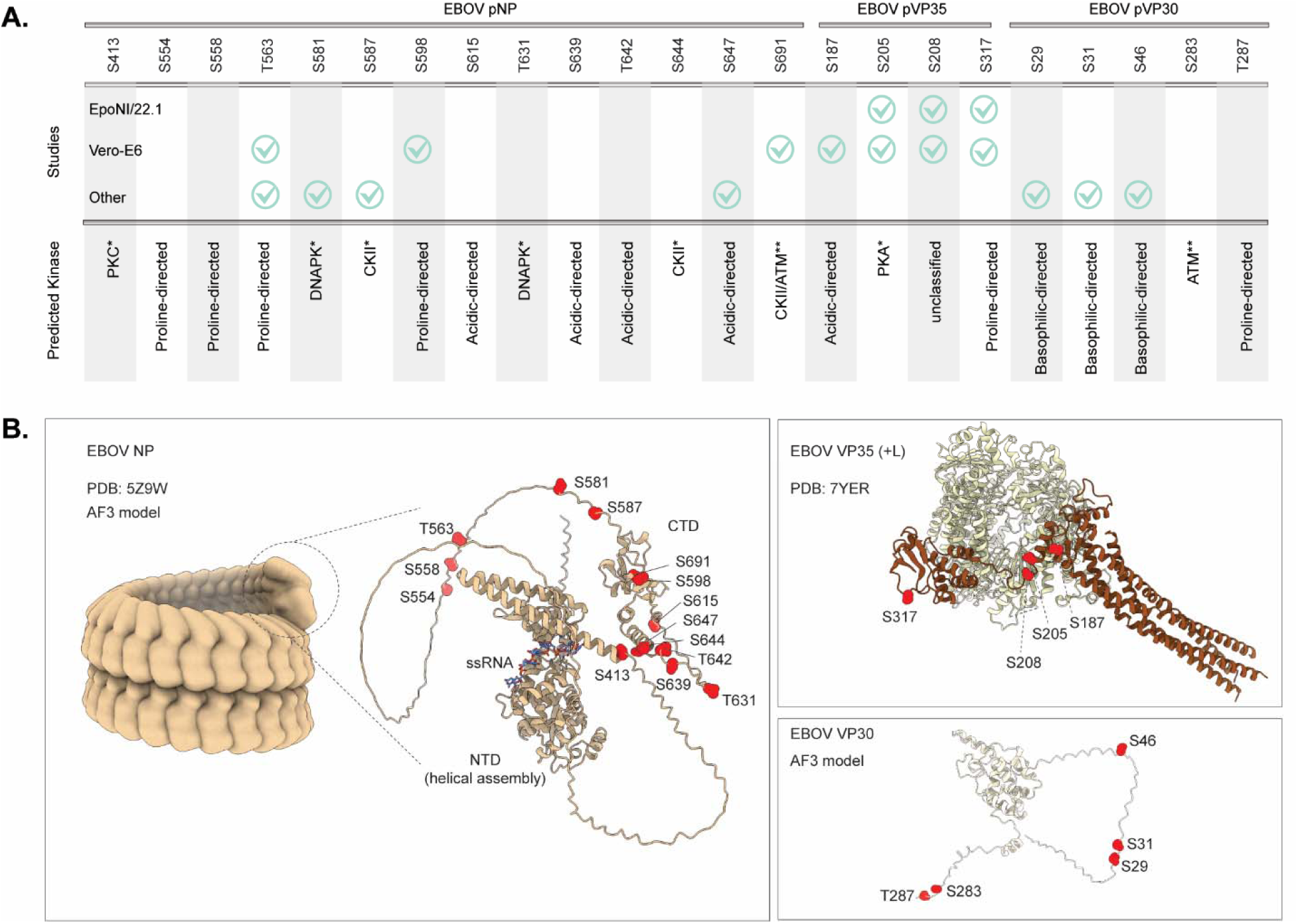
Cellular phosphorylation landscape of the EBOV RNP complex. **(A)** Localized detected phosphorylation sites (pSites) for NP, VP35 and VP30 proteins across all tested conditions (as depicted on the top). The pSites that have been previously identified in bat cells (EpoNI/22.1), kidney African green monkey cells (Vero-E6) and different human cell lines are check marked in mint green (25,39–41). On the bottom, the kinase prediction analysis for each individual EBOV protein phosphosite is depicted: * represents a kinase predicted by Netphos-3.1b (42); ** represents a kinase predicted by Arslan et al (43). For the pSites lacking kinase prediction, the kinase substrate sequence-preference motif was used for annotation. (B) Experimental and AlphaFold3 predicted models of EBOV RNP complex proteins (generated using PDB templates 5Z9W and 7YER) with pSites annotated.

The viral phosphorylation sites identified in our study showed various degrees of conservation across other *Orthoebolavirus* species (Figure S2). The detected phosphorylation sites of VP30 (S29, S31, S46, and S283) were found to be fully conserved across all species, with the exception of S287, situated in the far C-terminus of the protein (Figure 2B). In VP35, sites S205 and S317 are fully conserved across all species, while S187 and S208 carry potential phosphorylation acceptor sites only in different subsets of species. Interestingly, phosphomimetic mutations of S187 to D or A enhance or abolish EBOV replication, respectively, and species SUDV and RESTV both naturally carry the D187 substitution (39). In NP, phosphorylation sites S413, S587, and S691 are fully conserved, S563 and S581 are conserved in 5/6 species except for BOMV, and the remaining sites show a high degree of variability across species. The detected phosphorylation sites of NP all mapped outside of the structurally well characterized N-terminal domain that binds viral RNA and assembles into the tightly wound helical filaments characteristic of filoviruses (Figure 2B). Rather, the NP phosphorylation sites map to the extensive unstructured linker region between the N-terminal helical assembly and the functionally elusive C-terminal domain, as well as to the C-terminal domain itself. Likewise, VP35 phosphorylation sites map outside of the N-terminal oligomerization domains, but rather to the outer surface of the L-interacting C-terminal domain, all outside the L-VP35 interface. VP30 phosphorylation occurs mostly in the flexible N- and C-termini of this protein.

We compared these identified phosphorylation patterns of EBOV proteins expressed in HEK293T cells with those previously reported in human (HepG2, HEK293), monkey (Vero E-6) and bat (EpoNi/22.1) cell lines (Figure 2A) (25,39–41). The vast majority of VP35 phosphorylation sites were broadly conserved across all examined cell types, whereas VP30 phosphorylation sites were consistently observed in human cells but not detected in monkey or bat cells. NP phosphorylation displayed the most heterogeneous patterns across cell types and studies, with most sites identified in this work representing previously unreported phosphorylation events.

### EBOV minigenome activity remodels the host kinase signaling phosphoproteome

We focused next on how EBOV replication and transcription affects the host cell kinase signaling pathways, starting with phosphorylation dynamics of immune response related proteins. Among all significantly regulated phosphosites, 59 (18.5%) mapped to proteins annotated under ‘immunity’ or ‘viral infection’ in Uniprot, representing a substantial fraction of our dataset (Figure 3B). Several proteins with significantly regulated phosphorylation sites, including C1QBP and CD300A, were also annotated in the WikiPathway term “EBOV infection in host,” as noted above. CD300A functions as a negative regulator of TLR signaling, whereas C1QBP has been reported to translocate to mitochondria upon viral infection to inhibit innate immune signaling via the RIG-I/MAVS pathway. Notably, the C1QBP phosphorylation site S201 showed the largest fold change and is located within the mitochondrial targeting sequence. In addition, we also identified upregulation of phosphorylation site S599 of ADAR, a double-stranded RNA binding protein capable of editing viral RNA to increase translational efficiency of EBOV (44).

**Figure 3.**
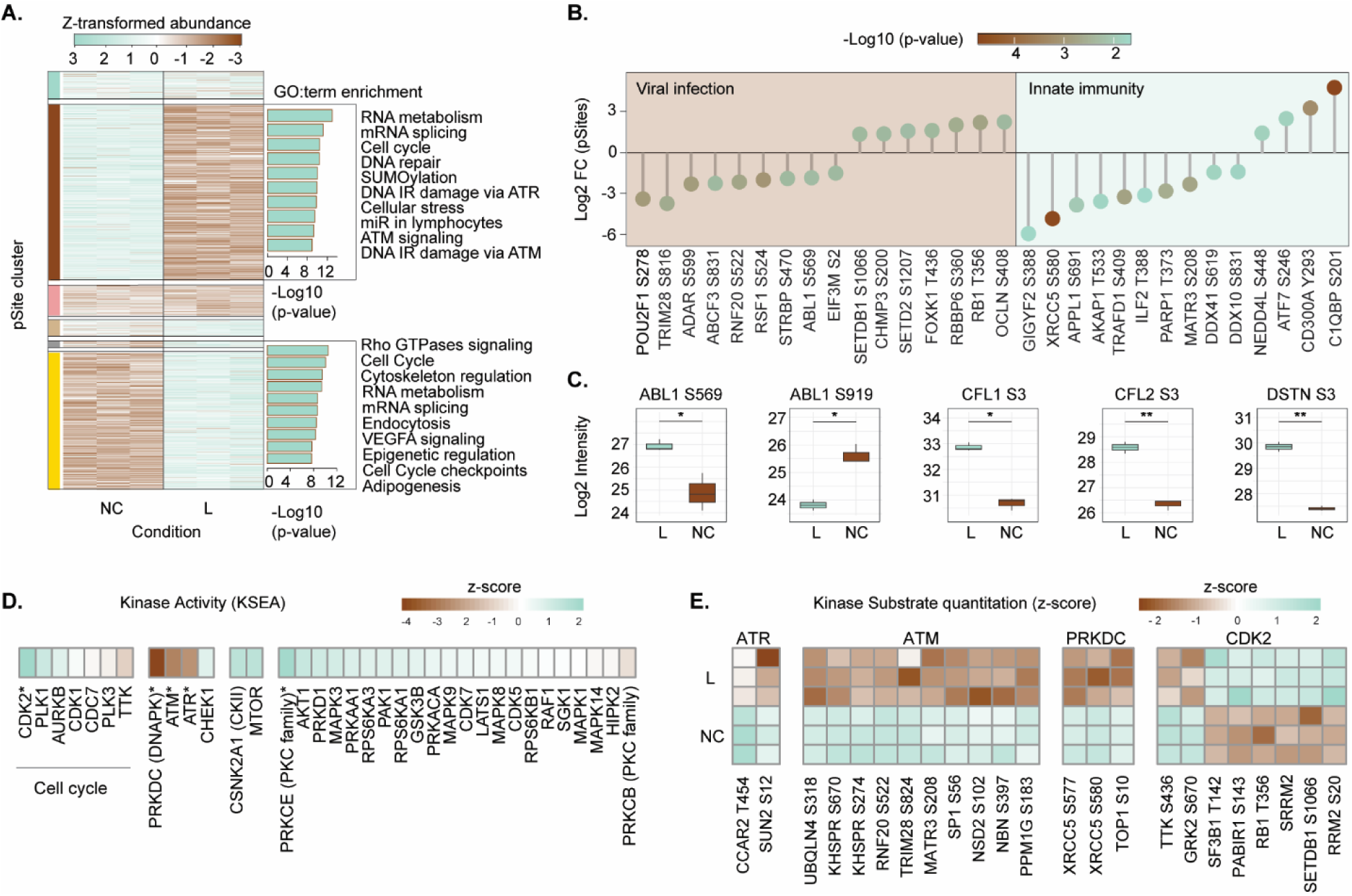
Host signaling rewiring in response to EBOV replication/transcription. **(A)** Hierarchical clustering of significantly changing pSites in L versus NC conditions is depicted on the left. Enriched non-redundant top 10 reactome and WikiPathways terms are depicted for each cluster (adjusted p-value < 0.001) on the right. Data were standardized using Z-scoring. **(B)** The lollipop plot depicts significantly regulated pSites of the proteins that previously have been shown to be involved in various viral infections (brown) or in dysregulation of innate immune response. Log2 fold change (FC) of pSites (L vs. NC) is shown. The color corresponds to p-value of the pSite FC change (adjusted p-value < 0.05). **(C)** Boxplots of Log2-transformed pSite intensities for proteins downstream of cytoskeletal rearrangement signaling. *: p-value < 0.05; **: p-value < 0.01. **(D)** Kinase–substrate enrichment analysis (KSEA), performed using the NETWORKIN algorithm on known substrates within Phosphomatics web tool, indicates increased activity of CDK2 and PRKCE and decreased activity of DNA damage response kinases (DNAPK, ATM, and ATR); *:p-value < 0.05. (45,46). **(E)** The heatmap of the kinase substrate quantitation for the kinases with significantly regulated activity.

Moving beyond the 67 phosphorylation sites annotated for ‘immunity’ and ‘viral infection’, hierarchical clustering of all significantly changing phosphorylation events identified two main clusters of differentially up- or downregulated sites between transfection control (NC) cells and EBOV minigenome replicating and transcribing cells (L) (Figure 3A, left). For both clusters, an enrichment analysis was performed to identify the pathways supporting EBOV minigenome activity. The cluster of downregulated phosphosites showed strong enrichment in the DNA damage response (DDR) pathway noted above (terms: DNA repair, DNA IR damage via ATR, ATM signaling and DNA IR damage via ATM). In contrast, the cluster of upregulated sites was primarily associated with cytoskeleton remodeling (terms: Rho GTPases signaling and cytoskeleton regulation). Indeed, STRING network analysis of host proteins harboring the phosphosites from the upregulated cluster revealed extensive involvement (∼15% of all proteins with regulated phosphosites) of cytoskeletal components including microtubule and actin networks as well as a Rho signaling node, interconnected with host proteins engaged in various viral processes (Figure S3B-C). Closer examination of the RHOA GTPase-mediated remodeling pathway, which includes the downstream kinases ROCK1/2 and LIMK1/2, revealed a significant increase in phosphorylation of downstream effector proteins directly involved in actin stress fiber formation, such as cofilin (CFL1 and CFL2) and destrin (DSTN) (Figure 3C) (47). We also observed differential phosphorylation patterns on ABL1, a key regulator of cytoplasmic mobility through actin filament reorganization (Figure 3C) (43,48). These findings point to the involvement of intermediate filaments as discussed in more detail below. Both up- and downregulated clusters contained phosphorylation sites involved in RNA metabolism, mRNA splicing and cell cycle control (terms: cell cycle and cell cycle checkpoints), highlighting disrupted cellular regulation and extensive kinase signaling rewiring in response to EBOV replication and transcription.

To further explore these perturbations, we inferred specific kinase activity from the experimentally determined phosphorylation patterns in our study, based on the known downstream substrates (Figure 3D) and phosphorylation site motif-based predictions (Figure S4A) (45,46,49). Both algorithms were applied to reduce bias, as not all phosphorylation sites can be assigned to a specific kinase and many kinases exhibit substrate promiscuity. From the subset of 35 kinases that showed signs of altered activity in EBOV replicating cells (Figure 3D, 3E, S4A), five exhibited the most striking changes beyond the statistical threshold used for this test (Figure 3D and S4B). CDK2 and PRKCE showed the strongest activation, whereas DDR pathway kinases (DNA-PK, ATM and ATR) displayed the most pronounced decrease in activity. The phosphorylation patterns of their identified substrates mirrored these activity trends, further supporting the kinase activity predictions (Figure 3E). Kinase–substrate analysis based on site motif predictions identified CDK6, CDK18, CDK2, and CDK4 among the five most active kinases (Figure S4B). These findings point to CDK2 and other cyclin dependent kinases as potential regulatory nodes during EBOV replication. Within the broader subset of 35 kinases with signs of altered activity, several were linked to cytoskeleton regulation (CKII, PAK1, CDK5), reinforcing the observation that cytoskeleton remodeling forms part of the broader cellular adjustments during EBOV replication. Furthermore, among aberrantly active kinases, several functional groups were distinguished including cell cycle regulators (CDK1/2/7, CDC7, PLK1/3, AURKB and TTK), cell growth-associated kinases (AKT1, MAPK1/3, RAF1, PRKACA) and DNA damage response kinases (DNA-PK, ATM, ATR and CHEK1). Hippo pathway kinases LATS1 and LATS2, previously implicated in EBOV virion egress, also showed a modest increase in activity (28).

### Host cell remodeling via cyclin dependent kinases is required for minigenome activity

Since CDK2 and other cyclin dependent dysregulated kinases were identified as potential regulatory nodes during EBOV replication, we hypothesized that their inhibition could suppress EBOV replication and transcription. To investigate this, we tested to what extent EBOV minigenome activity functionally relies on the identified kinase pathways by treating cells with various well-characterized and widely available small molecule kinase inhibitors. We tested inhibitors targeting the most strongly activated cyclin-dependent kinases (CDK2, CDK4/6, and CDK1) in our phosphoproteomics experiments, as well as other moderately activated signaling pathways. These include Hippo signaling (previously implicated in EBOV infection (28)); mTOR, AKT, and PKC pathways, which are known to be involved in other viral infections (50–52); and PAK1, a known regulator of cytoskeleton rearrangement (47). The inhibitors were chosen based on their specificity against each kinase. The PAK1 inhibitor did not have any notable effect on EBOV minigenome reporter signal (Figure S5A). Inhibitors of mTOR, AKT, PKC and Hippo signaling had only marginal effects on minigenome activity, even at the highest tested concentrations (10 µM) (Figure S5). In contrast, cyclin dependent kinase inhibitors potently diminished the eGFP reporter expression (Figure 4). This inhibitory effect was apparent both after inhibitor treatment 1 hrs prior to transfection and 4 hrs post transfection. Both CDK1 and CDK4/6 inhibitors decreased eGFP expression at higher tested concentration (ca. 10 µM) in the pre-treated cells only. In contrast, CDK2 inhibitors were particularly potent in a concentration-dependent manner, with reduced eGFP expression observed even at lower concentrations (0.5 and 2.5 µM), independent of treatment time. The cyclin dependent kinase inhibitors had no visible effects on cell viability at the tested concentrations (Figure 4A), indicating that their effect on EBOV minigenome activity should be attributed specifically to their action on the targeted kinases. Differential inhibition of eGFP reporter expression by CDK inhibitors targeting distinct cell cycle stages suggests that EBOV replication and transcription modulate the cell cycle towards an S/G2-like state. Consistently, these experiments indicate a functional reliance on cyclin-dependent kinase signaling, particularly CDK2, to sustain viral replication and transcription.

**Figure 4.**
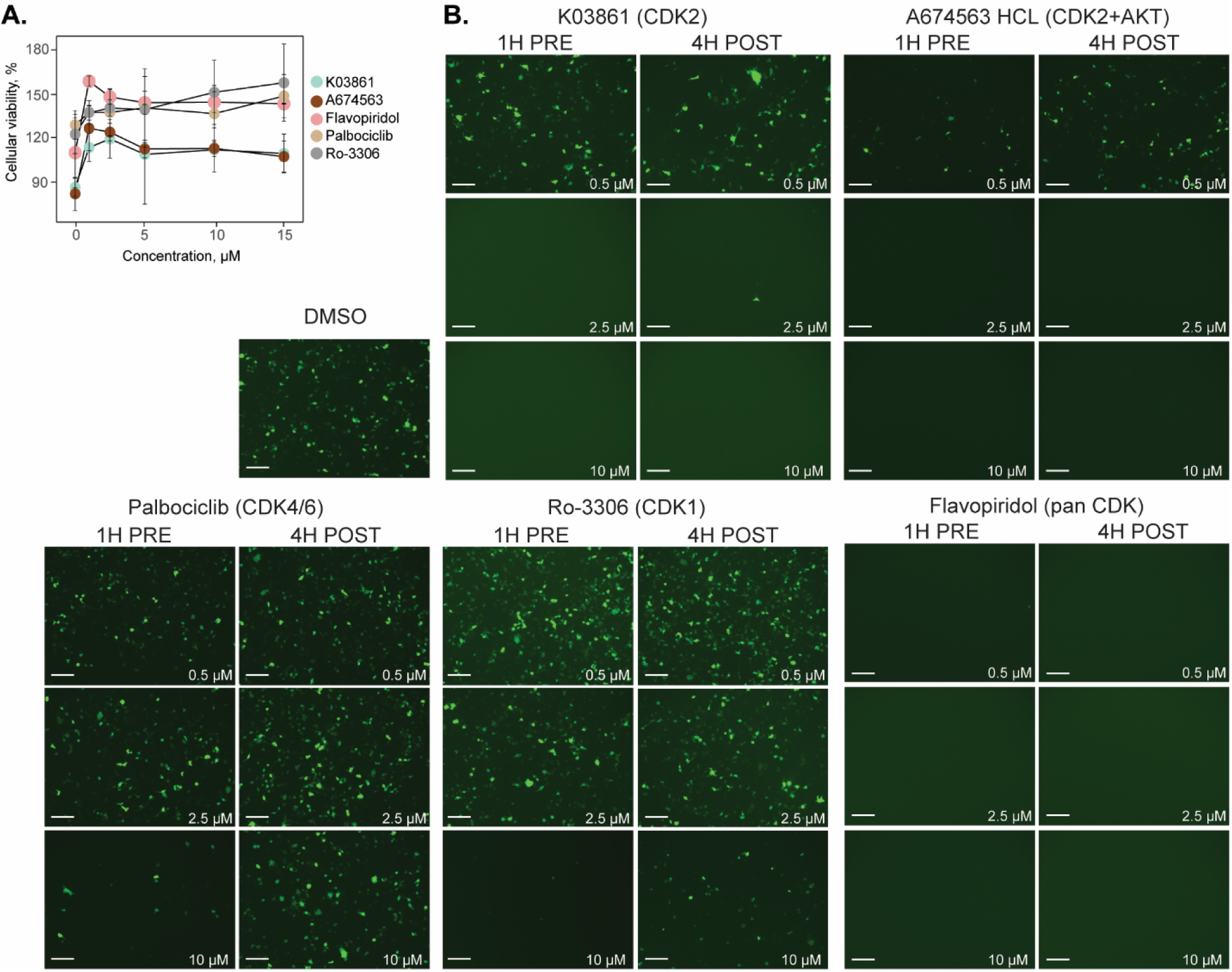
Inhibition of CDK2 and other cyclin dependent kinases blocks EBOV transcription/replication. **(A)** Cell viability of cells treated with CDK inhibitors across a range of concentrations, relative to DMSO control. **(B)** Representative images of the HEK293T cell cultures treated with CDK2 (K03861; A674563), CDK4/6 (Palbociclib), CDK1 (Ro-3306) and panCDK (Flavopiridol) inhibitors or DMSO 1 H before or 4 H after the minigenome system transfection for 18 hours. The GFP signal represents cells with an active EBOV replication/transcription complex. Kinase inhibitor concentrations used for treatment are indicated in the lower right corner (0.5, 2.5, and 10 µM). Scale bars, 300 µm.

## Discussion

Using (phospho-)proteomics profiling of the host cell response to EBOV minigenome activity, we provide a systems wide view at the host cell remodeling required to facilitate ebolavirus replication and transcription. Our findings show that minigenome activity is associated with a moderate shift in the host proteome, with fewer than 40 proteins exhibiting differential expression in the presence of actively replicating and transcribing ebolavirus RNP’s. Amongst the significantly changing proteins, TRIM14, ZDHHC20, SOX2, FLCN, and ATXN1L have been previously implicated in the host cell response to EBOV and other viruses (31,33–35,37). Overexpression of TRIM14 was found to suppress EBOV replication, interact with EBOV-NP and promote interferon-β and NF-κB activation (31). Moreover, ZDHHC20 is known to palmitoylate interferon-induced transmembrane protein 3 (IFITM3) to enhance its antiviral activity (32,37). SOX2 transcription has been shown to be upregulated in HUVECs following EBOV infection (33). Upregulated FLCN transcription has been shown to be a predictor of poor vaccine responses for EBOV immunization and is linked to mTORC1 related signaling (34). ATXN1L is a transcriptional repressor of IFN and ISG via a Capicua (CIC) related mechanism (35). Notably, we also observed minigenome-induced overexpression of CKS2 and WTIP. CKS2 is a subunit of cyclin-dependent kinases, which exhibited some of the strongest signs of altered activity in our phosphoproteomics results. Likewise, WTIP is an adaptor protein in Hippo signaling (36) which also showed increased activity in our phosphoproteomics analysis (LATS1 and LATS2 kinase activity) and was previously linked to the regulation of EBOV transcription and egress (28).

Although these changes in the proteome were significant, they were moderate in amplitude. In contrast, we observed major changes in the phosphoproteome, with over 300 differentially regulated phosphorylation sites in response to minigenome activity in HEK293T cells, including 23 sites on the ebolavirus proteins. A notable subset of the viral phosphorylation sites is linked to DNA damage response (DDR) pathways, in line with altered DDR kinase activity observed across host protein phosphorylation. A previous bioinformatic study identified more potential ATM kinase recognition motifs on all EBOV proteins (43), while another study reported co-localization of ATM and ATR kinases to the EBOV replication factories in infected cells (53). Furthermore, known substrates of the DDR kinases (ATM, ATR, and DNAPK) showed reduced phosphorylation in our study, whereas a previous study has reported activation of IFN responses upon small molecule induction of ATM signaling and engagement of the DNA sensor cGAS and its adaptor STING (54). Collectively, these results highlight a critical role for DDR-associated phosphorylation in EBOV replication, potentially involving sequestration of DDR kinases into viral replication factories and suppression of IFN signaling via ATM and cGAS–STING pathways. Exactly how DNA damage is involved in EBOV replication is currently not clear, but it is likely that cellular stress related to viral replication may induce DNA damage and concomitant DDR pathway stimulation via ATM, ATR and cGAS-STING signaling. It may also be the case that EBOV replication triggers DDR pathways via DNA damage-independent mechanisms and crosstalk via other signaling pathways. Notably, other viruses, including hepatitis C virus and SARS-CoV-2, have also been shown to modulate DNA damage response (DDR) pathways to support efficient replication (37,55–57).

The changes observed in host protein phosphorylation point to a central role for DDR, cell cycle control, immune response regulation and cytoskeletal remodeling in facilitating ebolavirus minigenome activity. Cytoskeletal remodeling appears to follow a RHOA GTPase-mediated pathway, including downstream kinases ROCK1/2, LIMK1/2, PAK1 and effector proteins directly involved in actin stress fiber formation, such as cofilin (CFL1 and CFL2) and destrin (DSTN)(47). We also observed differential phosphorylation patterns on ABL1, a key regulator of the cytoplasmic mobility through actin filament reorganization (43,48). Consistent with this, functional EBOV viral factories have been shown to depend on physical interactions with intermediate filaments (58). Studies further show that intermediate filaments in turn rely on physical association with actin filaments to form composite cytoskeletal networks and regulate the excessive stress fiber formation (58,59). Together, these findings suggest that EBOV RNP components, potentially through unbound VP35, hijack cytoskeleton regulation pathways during EBOV replication to facilitate the proper formation and stability of viral factories.

With respect to immune regulation, many significantly regulated phosphorylation sites mapped to proteins known to be involved in viral infection and immune response pathways. In fact, the strongest observed change occurred for C1QBP S201, within the mitochondrial targeting sequence, raising the possibility that phosphorylation at this site may contribute to regulation of mitochondrial translocation and associated immune signaling (60). Whether and how this phosphorylation of the mitochondrial translocation sequence of C1QBP plays a role in RIG-I/MDA5 signaling via MAVS is yet to be determined.

Testing a variety of kinase inhibitors revealed that EBOV minigenome activity relies strongly on cyclin dependent kinases. The pattern of cell cycle kinase activity observed in our experiments is consistent with an S/G2-like state, characterized by increased CDK2 activity and reduced activity of CDK7 and CDC7 (G1/S transition), along with decreased EBOV minigenome eGFP reporter signal in cells pre-treated with CDK4/6 and CDK1 inhibitors, which arrest cells in G1 and S phases, respectively (61). Furthermore, decreased activity of PLK1 (G2/M), together with reduced TTK and PLK3 activity, is indicative of suppressed mitotic signaling (62,63). These findings add to a growing body of evidence that cell cycle regulation is an important aspect of host cell remodeling to facilitate viral replication and transcription across diverse families of viruses (64–67). The functional importance of cell cycle regulation via cyclin dependent kinases points to a possible pathway for the development of a host-directed antiviral strategy against *Orthoebolaviruses* as inhibition of these pathways not only suppressed EBOV replication machinery in our study but has also been shown to impair viral replication in infected cells in a previous study (68).

Collectively, our data indicate that EBOV induces extensive host cell remodeling characterized by suppression of DDR signaling, and activation of PKC, AKT, mTOR, Hippo, and especially CDK2-dependent pathways. These findings point to innate immune suppression, cytoskeletal remodeling, and cell cycle regulation as central drivers of EBOV replication and transcription.

## Materials and methods

### Minigenome plasmids

The Ebola minigenome assay plasmids were purchased from Addgene: pCAGGS_3E5E_eGFP (ID: 103054); pCAGGS_L_EBOV (ID: 103052); pCAGGS_VP30_EBOV (ID: 103051); pCAGGS_VP35_EBOV (ID: 103050); pCAGGS_NP_EBOV (ID: 103049) (15). Experiments were performed at containment level ML-II (BSL-2), following The Netherlands Commission on Genetic Modification (COGEM) advice CGM/210729-01.

### Cell culture

HEK293T cells were cultured in Dulbecco’s modified Eagle’s medium (DMEM) supplemented with (or without) 10% fetal bovine serum (Sigma) in a humidified incubator set to 5% CO2 at 37°C.

### Minigenome assay

HEK293T cells were seeded in 10-cm dishes with 2.4 million cells in FBS-free medium and cultured for 24 hours. After 24 hours, medium was switched to 10% FBS medium and cultured for another 24 hours. Next, each plate of cells was transfected with total of 15 µg of DNA of the minigenome assay plasmids (L, NP, VP35, VP30 and minigenome(MG)) using TransIT (Merus) transfecting agent according to the manufacturer’s protocol. The following plasmid ratio’s were used NP: VP35: VP30: L: MG = 2.5: 2.5: 1: 40: 15 (w/w). The transfected cells were grown for either 6, 24 or 48 hours depending on the subsequent analysis. The minigenome transfection efficiency was assessed by the intensity and relative count of GFP producing cells.

### Sample preparation for proteomics

Cells were collected either 6 or 24 hours post transfection as follows: medium was removed, cells were washed with PBS and collected by scraping with 5 ml of ice-cold PBS, centrifuged at 300 g for 10 minutes, followed by aspiration of PBS. The collected cells were frozen in liquid nitrogen. Each frozen cell pellet was resuspended in 750 µl of SDC lysis buffer (100LmM Tris pH 8.5, 1% sodium deoxycholate, 10 mM tris(2-carboxyethyl)phosphine, 40LmM chloroacetamide) supplemented with phosphotase inhibitor tablet (PhosSTOP, Roche) and protease inhibitor cocktail tablet (cOmplete mini EDTA-free, Roche) and sonicated with a a Bioruptor Plus (Diagenode) for 15 cycles of 30 s. Then, lysates were denatured at 95 °C for 5 minutes with the follow-up reduction and alkylation for 20 minutes in the dark. Each sample was diluted in 1:10 ratio (v/v) with 50 mM ammoniumbicarbonate, pH 8.0. A BCA protein assay was used to determine the concentration of the acquired protein content. 200 µg of protein were digested overnight with trypsin (Sigma) and lysyl endopeptidase (Lys-C, Wako) in 1:50 and 1:75 ratio of enzyme:protein (w/w) respectively at 37 °C. SDC was precipitated with formic acid at 2% (final concentration) and samples were centrifuged at 21,100L×L*g* for 20 min at 4 °C. Next, acquired peptides were desalted with 10 mg Oasis HLB 96-well plate (Waters) as previously described (69) and vacuum-dried.

### Phosphorylated peptide enrichment

Phosphorylated peptides were enriched from 200 µg peptide input in the automated fashion with the AssayMAP Bravo Platform (Agilent Technologies) according to the published protocol by Post et al (70).

### LC-MS/MS for proteomics

The peptides were dissolved in 2% formic acid to the concentration of 1 μg/μl. One μl of resuspended peptides was analyzed on an Orbitrap Exploris 480 (ThermoFisher Scientific, Bremen) coupled online to UHPLC system Ultimate 3000 (ThermoFisher). Peptides were loaded onto an Acclaim Pepmap 100 C18 (5LmmL×L0.3Lmm, 5 μm, ThermoFisher Scientific) trap column and separated with in-house made 50-cm reverse-phase analytical column (75 μm inner diameter, ReproSil-Pur C18-AQ 2.4 μm resin [Dr. Maisch GmbH]) with a 120-min gradient starting at 9% buffer B to 13% B in 1 min, from 13% to 44% in 96 min, from 44% to 55% in 6Lmin, from 55% to 99% in 1 min, 99% wash-out in 5 min and re-equilibration back to 9% buffer B in 10 min where buffer A is 0.1% formic acid in water and buffer B is 0.1% formic acid in 80% acetonitrile (v/v). The data were acquired in data-independent mode. Exploris 480 parameters for the full MS scans were as follows: scan mass range set to 375-1600 *m/z*, resolution of 60 000 at 200 *m/z*, AGC target set to standard, maximum injection time set to auto. For the MS/MS events the following parameters were used: cycle time of 3s, quadrupole isolation window of 20 Da with 1 Da overlap between the windows (total of 30 windows), the precursor mass range was set to 400-1000 *m/z*, resolution of 30 000 at 200 *m/z*, normalized AGC target was set to 1000%, injection time set to auto, normalized collisional energy was set to 28%.

### LC-MS/MS for phosphoproteomics

The phosphopeptides were resuspended in 11 µl of 20 mM citric acid in 2% formic acid. Five μl of resuspended peptides were analyzed Exploris 480 (Thermo Scientific, Bremen) coupled online to UHPLC system Ultimate 3000 (ThermoFisher). Peptides were loaded onto an Acclaim Pepmap 100 C18 (5LmmL×L0.3Lmm, 5 μm, ThermoFisher Scientific) trap column and separated with in-house made 50-cm reverse-phase analytical column (75 μm inner diameter, ReproSil-Pur C18-AQ 2.4 μm resin [Dr. Maisch GmbH]) with a 120-min gradient starting at 9% buffer B to 36% B in 98 min, from 36% to 55% in 6 min, from 55% to 99% in 1 min, 99% wash-out in 5 min and re-equilibration back to 9% buffer B in 10 min where buffer A is 0.1% formic acid in water and buffer B is 0.1% formic acid in 80% acetonitrile (v/v). The data were acquired in data-dependent mode. Exploris 480 parameters for the full MS scans were as follows: an AGC target of 5L×L10^4^ at 60 000 resolution, scan range 375-1600L*m/z*, Orbitrap maximum injection time set to auto. The MS2 spectra were acquired for the ions in charge states 2+ to 6+, at a resolution of 30,000 with an AGC target of 5L×L10^5^, maximum injection time set to auto, scan range was set from the first mass of 120L*m/z*, and dynamic exclusion of 16Ls, precursor ion selection was at 1.4L*m/z*. The fragmentation was induced by stepped higher collision-induced dissociation with the NCE set to 15, 28, 35.

### (Phospho)proteomics data processing and analyses

For the proteomics data, raw files were searched with DIANN (version 1.8.1) with library free search enabled and for the phosphoproteomics data (71), files were searched with MaxQuant (version 2.1.0.0) (72) against Uniprot human database supplemented with the protein sequences of Zaire ebola virus (version March 17, 2022). Carbamidomethylation of cysteine was specified as a fixed modification, whereas oxidation of methionine, and phosphorylation of serine, threonine, and tyrosine were included as variable modifications. Trypsin was set as enzyme, allowing up to two missed cleavages. The filtering was performed at 1% false discovery rate (FDR) at protein and peptide level. Quantitative data were processed using Perseus (version 1.6.14.0) to remove contaminants and reverse hits, followed by filtering for localization probability >0.75 (phosphoproteomics data). Data were log2-transformed and median-normalized across columns. Matrices were further filtered for data completeness, requiring at least two values per group.

Downstream analyses, including principal component analysis (PCA), significance testing and clustering, were performed on the processed dataset. Phosphorylation and kinase-substrate enrichment analyses were performed using Phosphomatics (45) and Rokai (49) platforms. Gene ontology enrichment analyses were performed using Metascape (73) and String DB resources (74). Downstream analyses and visualization were performed in R using the following packages: Limma, ggplot2, Bioconductor, eulerr, gplots, pheatmap, tidyverse.

### Mapping phosphorylation of EBOV proteins onto structure

Experimentally determined phosphorylation sites were mapped to publicly available EBOV structures of NP, VP35-L, and VP30, or AlphaFold3 generated models. Structures were visualized with ChimeraX v1.9.

### Cell viability assay

To determine the cell viability under the kinase inhibitor treatment, cells were seeded in 96-well plates, transfected and treated with kinase inhibitors at concentrations of 0 µM, 0.5 µM, 2.5 µM, 5 µM, 10 µM and 15 µM. Next, 10 µl of AlamarBlue solution (Thermo Fisher Scientific) was added to the cells 6 hours post-transfection and incubated for 18 hours. The color change was measured using absorbance at 570 and 595 nm (plate reader Multiscan FC, Thermo Fisher scientific). The viability was calculated as percentage of live minigenome replicating cells compared to the DMSO treated control cells.

### Inhibition of kinase activity in EBOV minigenome replicating cells

HEK293T cells (65 000 cells/well) were seeded in 24-well plate in the cells in FBS-free medium and cultured for 24 hours. After 24 hours, medium was switched to 10% FBS medium and cultured for another 24 hours. Next, each well was transfected with minigenome assay plasmids as described above. The transfected cells were treated with kinase inhibitors at 0.5 µM, 2.5 µM or 10 µM concentration either 1 hour pre-transfection or 4 hours post-transfection.

### Live cell imaging

To assess the transfection efficiency of minigenome plasmids and the efficacy of kinase inhibitors in EBOV minigenome replicating cells, the corresponding cells were imaged 24 hours post-transfection with EVOS M5000 Imaging system (Invitrogen by ThermoFisher Scientific) equipped with GFP light cube and 10x phase objective. The images were taken with the dedicated EVOS software.

## Data availability

The mass spectrometry based proteomics and phosphoproteomics raw data were deposited on PRIDE under the dataset identifier PXD076120.

## Supplementary information

The supplementary information file contains Supplementary Figures S1-S5, Supplementary Table S1, and Supplementary Data S1.

## Acknowledgements

This research was funded by the Dutch Research Council NWO Gravitation 2013 BOO, Institute for Chemical Immunology (ICI; 024.002.009) to J.S. We would like to thank dr. P. van Breugel for creating the catalytically dead mutant of Ebola L protein, and dr. P. Sinitcyn for help with the code to survey the Uniprot function and GO terms.

## Declaration of possible conflicts of interests

All authors declare no competing interests.

