## Supplementary Information for "Host cell remodeling via cyclin dependent kinases drives Ebola virus replication and transcription"

- Supplementary Figures S1-S5
- Supplementary Table S1

**Supplementary Table S1. Netphos-3.1b** (1) **probability scores for the predicted host kinases to EBOV phosphorylation sites.** Scores >0.500 for a specific kinase indicate that the kinase is likely to target the phosphorylation site. The higher the score, the higher the probability the phosphosite of interest is targeted by the predicted kinase.

| pSite | EBOV protein | Netphos-3,1b score | Predicted kinase |
|---|---|---|---|
| S205 | VP35 | 0.547 | PKA |
| S644 | NP | 0.570 | CKII |
| T631 | NP | 0.633 | DNAPK |
| S587 | NP | 0.737 | CKII |
| S581 | NP | 0.567 | DNAPK |
| S413 | NP | 0.855 | PKC |

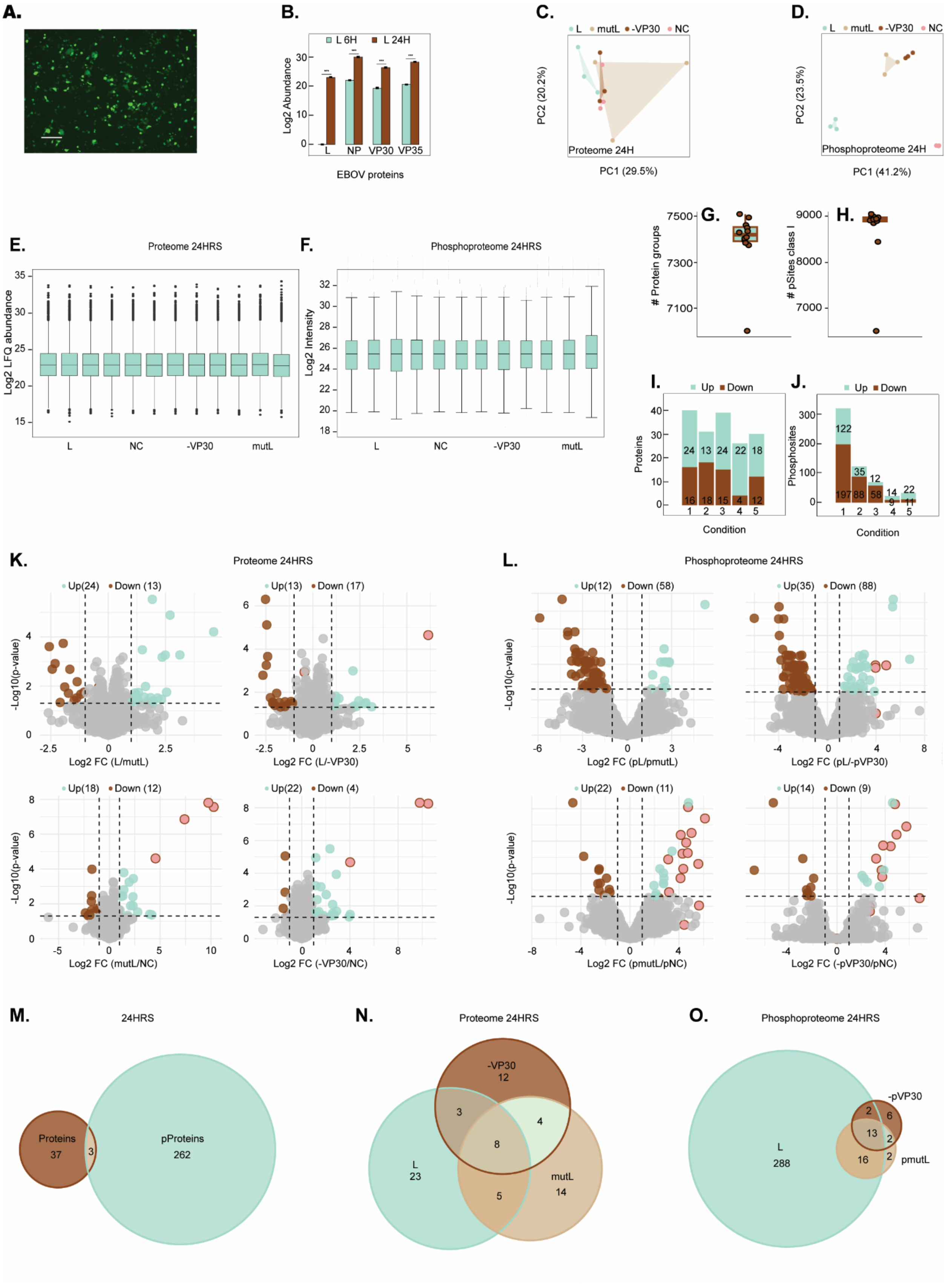
A.
B.
L 6H
L 24H
Log2 Abundance
EBOV proteins
L
NP
VP30
VP35
C.
L
mutL
-VP30
NC
PC2 (20.2%)
PC1 (29.5%)
Proteome 24H
D.
PC2 (23.5%)
PC1 (41.2%)
Phosphoproteome 24H
E.
Proteome 24HRS
Log2 LFQ abundance
F.
Phosphoproteome 24HRS
Log2 Intensity
G.
### Protein groups
H.
### pSites class I
I.
Up
Down
Proteins
Condition
J.
Phosphosites
K.
L.
-Log10(p-value)
Log2 FC (L/mutL)
Log2 FC (L/-VP30)
Log2 FC (mutL/NC)
Log2 FC (-VP30/NC)
Log2 FC (pL/pmutL)
Log2 FC (pL/-pVP30)
Log2 FC (pmutL/pNC)
Log2 FC (-pVP30/pNC)
M.
24HRS
Proteins
37
3
pProteins
262
N.
Proteome 24HRS
-VP30
12
L
23
mutL
14
O.
Phosphoproteome 24HRS
-pVP30
pmutL
L
288

**Figure S1. Overview of quantitative (phospho)proteomics data and quality control metrics. (A)** Representative image of active Ebola minigenome system 24 H post transfection. Transfection with EBOV L, VP30, VP35, NP, and minigenome (with eGFP reporter) plasmids leads to eGFP expression within 24 hours, indicative of active minigenome replication and transcription Scale bar, 300 µm. **(B)** Box plot of Log2-transformed protein abundances (LFQ; label free quantitation abundance). **(C)** Box plot of Log2-transformed phosphopeptide intensities. **(D)** Boxplot showing number of quantified protein groups. **(E)** Boxplot showing number of class I phosphosites used for quantification after filtering. **(F)** Principal-component analysis (PCA) of proteome and **(G)** phosphoproteome, with 3 replicates, 24 H post-transfection. The phosphoproteomes are strongly separated in both PC1 and PC2 components, the proteomes are less divergent. **(H)** Abundance of individual EBOV proteins 6 H and 24 H post-transfection. The L-protein was not detected 6 H post-transfection. The abundance of other RNP proteins went up at 24 H post-transfection. Due to the incomplete RNP complex formation after 6 H post-transfection, the subsequent analyses were performed on the 24 H post-transfection data. The *** represents a p-value <0.001. The bars represent the standard deviation. **(I)** Number of differentially abundant protein groups **(J)** and differentially abundant phosphosites across all conditions. Mint green represents the protein groups or phosphosites significantly increased in abundance; Brown represents the protein groups or phosphosites significantly decreased in abundance. Conditions: 1 (L/NC); 2 (L/-VP30); 3 (L/mutL); 4 (-VP30/NC); 5 (mutL/NC). **(K)** Volcano plot depicting Log2-transformed abundance fold change 24 H post-transfection for protein groups and **(L)** phosphosites for the following conditions: L over mutL; L over -VP30; mutL over NC and -VP30 over NC). Mint green represents increase in abundance (F C> 1, p-value <0.05). Brown represents the decrease in abundance (FC < -1, p-value <0.05). Pink highlights EBOV proteins or phosphosites. **(M)** Euler diagram shows the overlap between the significantly regulated protein groups (brown) and proteins with the regulated phosphosites (mint green). All 3 overlapping proteins were found to be EBOV proteins. **(N)** Euler diagram represents the overlap between significantly regulated proteins groups against the corresponding NC in the groups: L/NC; mutL./NC and -VP30/NC. **(O)** Euler diagram represents the overlap between significantly regulated phosphosites against the corresponding NC in the groups: L/NC; mutL./NC and -VP30/NC.

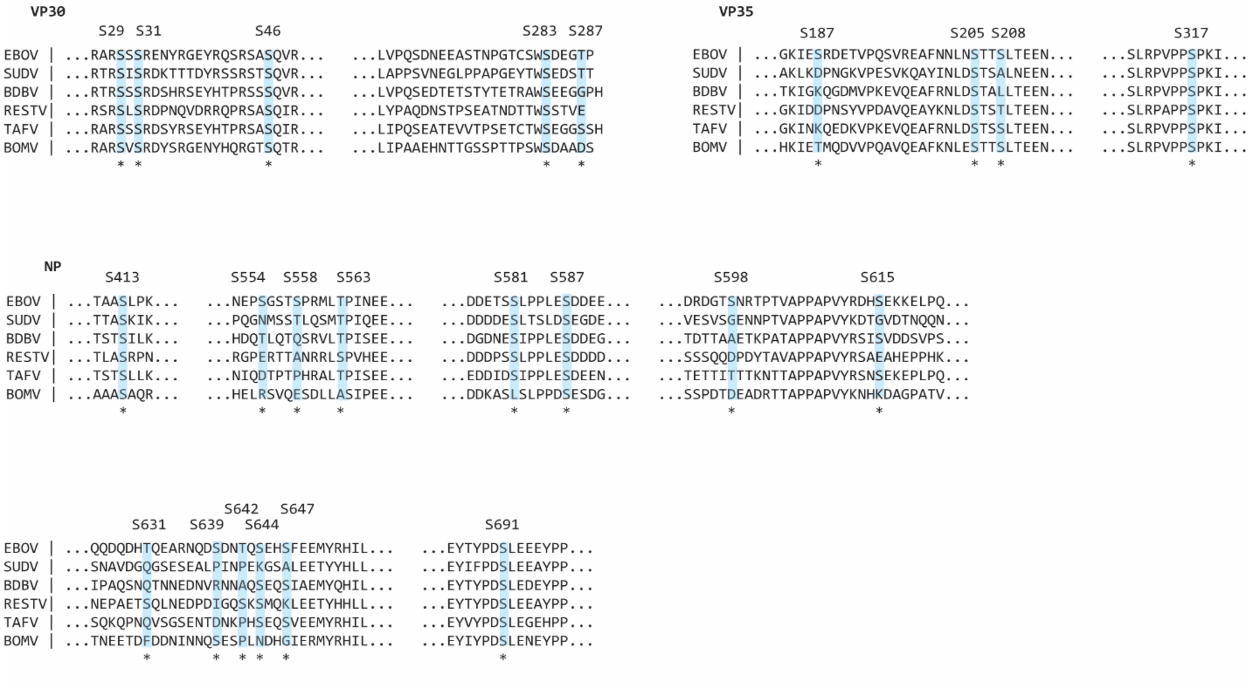


**Figure S2. Sequence alignment of the EBOV phosphorylated proteins (NP; VP35 and VP30) across different filoviruses.** The identified phosphorylation sites are highlighted with an asterisk. Only those parts of the sequences harboring detected phosphosites are shown.

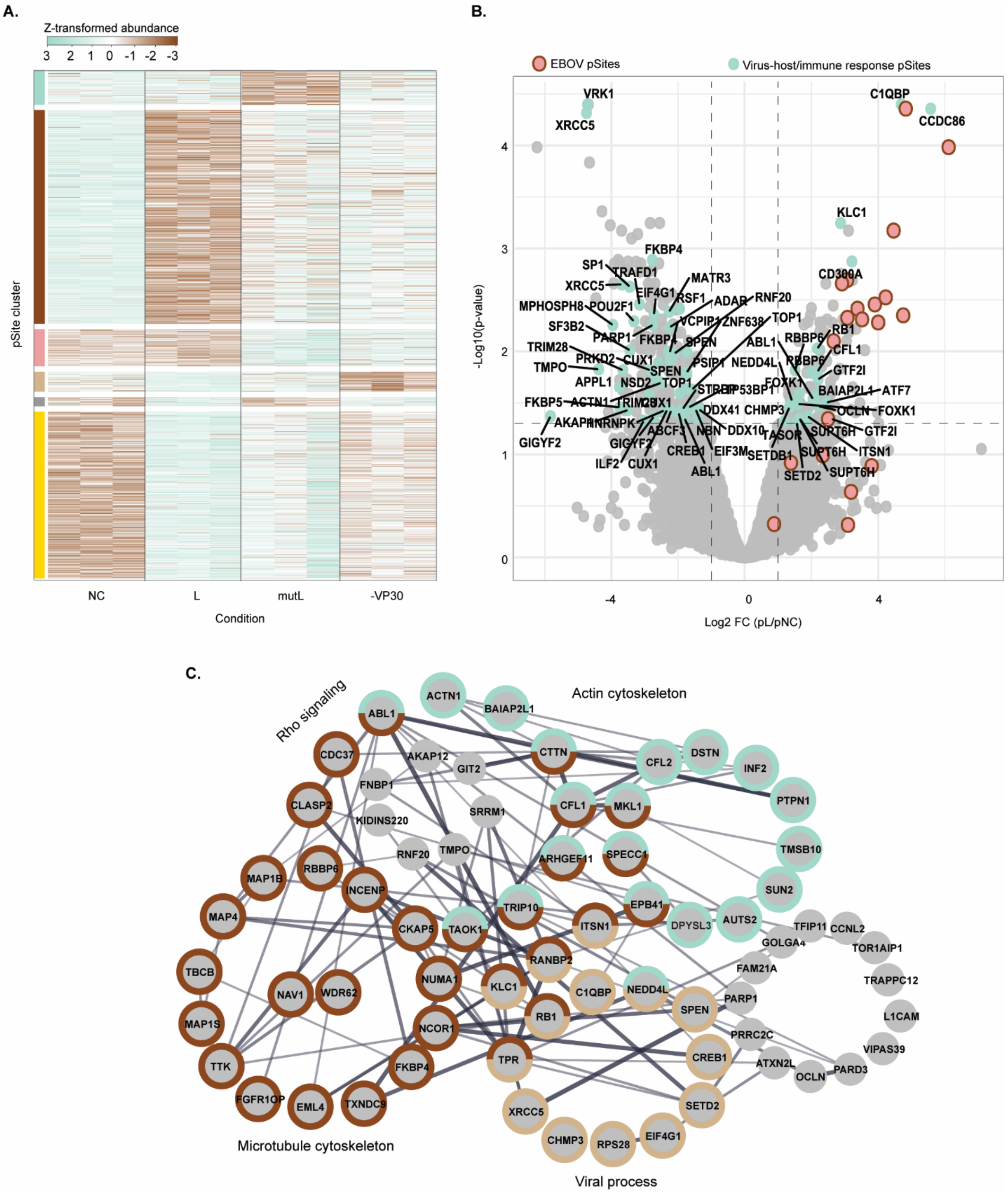


**Figure S3. Rewiring of host signaling networks and regulated phosphorylation sites. (A)** Clusters of significantly changing phosphosites under all tested conditions 24 H post-transfection. The full rewiring can be observed in L versus NC conditions. In contrast, mutL and -VP30 show milder changes in comparison to NC. **(B)** Volcano plot of log2-transformed abundance fold change (FC) of L condition versus NC 24 H post-transfection. Mint green represents all the differentially regulated phosphosites of host proteins involved in known host-virus interactions, virus infection or immune response regulation: a total of 59 out of 319 differentially regulated phosphosites (18.5%). **(C)** STRING network analysis of the host proteins with the regulated phosphosites. The colors of circles represent the process description (FDR value < 0.01). A substantial fraction of regulated phosphosites mapped to proteins involved in cytoskeletal organization, with 39 proteins identified in the STRING network out of 262 proteins with regulated phosphosites (15%).

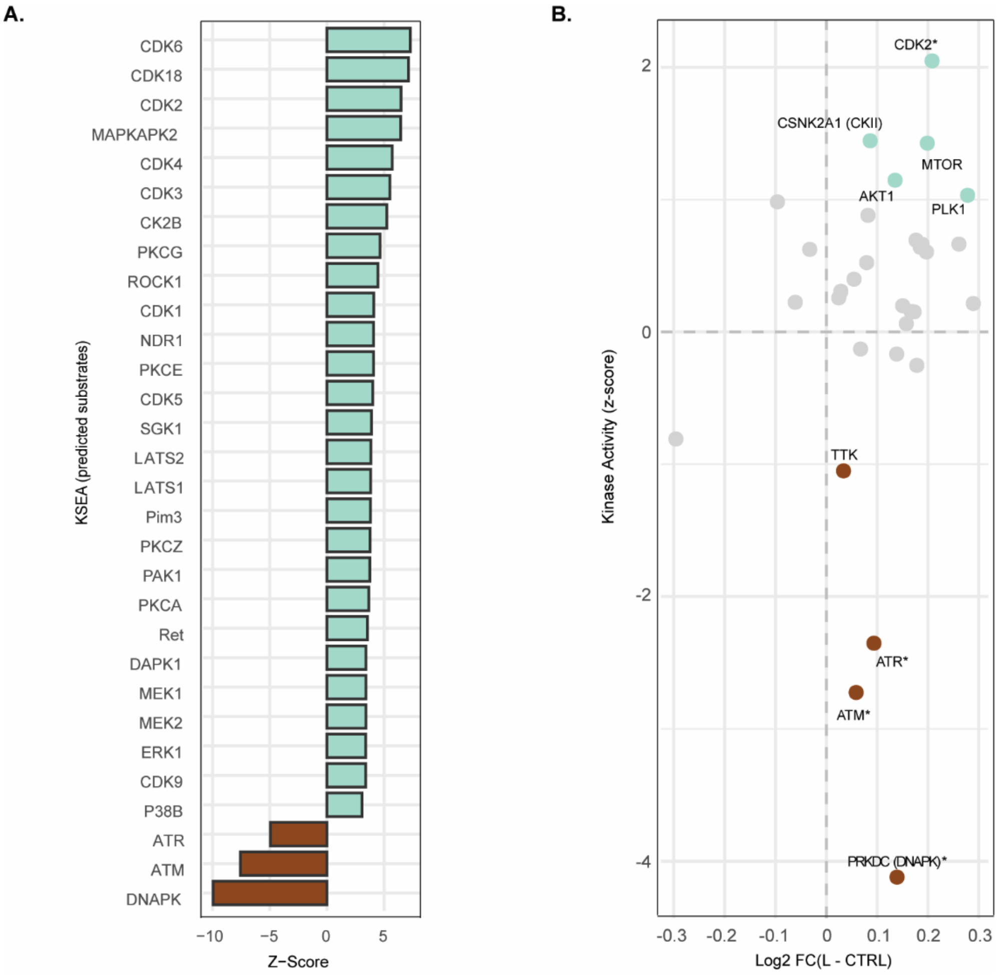


**Figure S4. Kinase abundance *versus* kinase activity based on Kinase-Substrate Enrichment Analysis. (A)** Kinase-Substrate Enrichment Analysis (KSEA) by ROKAI algorithm based on the phosphosites motif predicted substrates reveals a broader range of kinases with increased activity, with CDK6 showing the strongest activation. The kinases with decreased activities are based upon the NETWORKIN analysis with known substrates. **(B)** Log2-transformed fold changes in kinase abundance (x-axis) are plotted against corresponding changes in kinase activity (y-axis) comparing the L condition with the control. Clearly, activities of the kinases of interest don't depend directly on the kinase abundance in the cell.

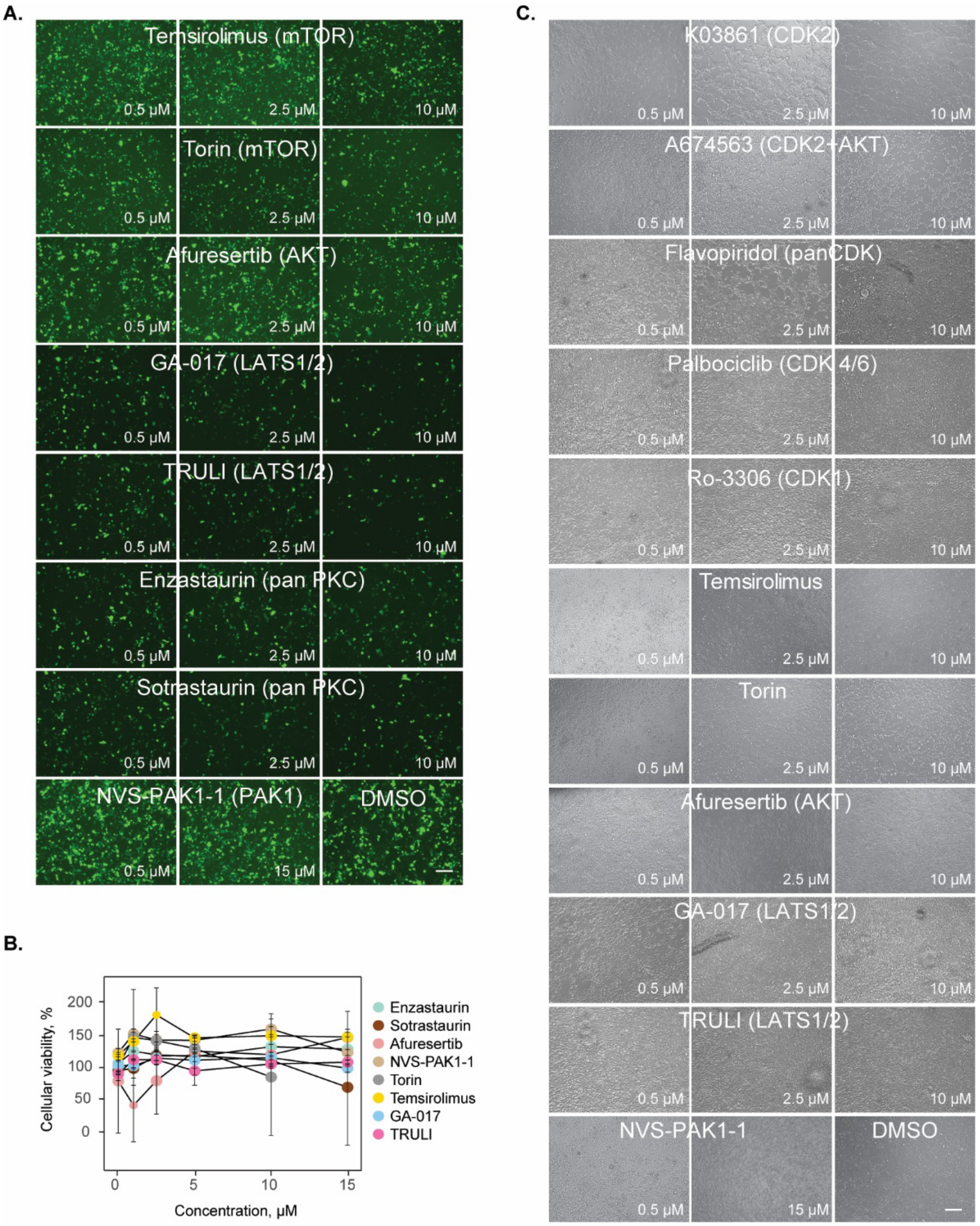


**Figure S5. Representative images visualizing the effect of the selected kinase inhibitors on the cells, actively replicating/transcribing the EBOV minigenome system. (A)** Representative images of HEK293T cell cultures treated with inhibitors targeting mTOR (Temsirolimus, Torin), AKT (Afuresertib), Hippo (GA-017, TRULI), panPKC (Enzastaurin, Sotrastaurin) and PAK1 (NVS-PAK1-1) or (as control) DMSO vehicle 4 H after the minigenome system transfection. The GFP signal represents cells with an active EBOV replication/transcription complex. Kinase inhibitor concentrations used for treatment are indicated in the lower right corner (0.5, 2.5, and 10 μM). Scale bars, 300 μm. **(B)** Cell viability determined for cells treated with the kinase inhibitors at various concentrations compared to a DMSO control. **(C)** Trans images of the cells treated with all kinase inhibitors in various concentrations in this study. All cells appear viable, albeit with aberrant phenotypes. Kinase inhibitor concentrations used for treatment are indicated in the lower right corner (0.5, 2.5, and 10 μM). Scale bars, 300 μm.
